# DeSciDe: A tool for unbiased literature searching and gene list curation unveils a new role for the acidic patch mutation H2A E92K

**DOI:** 10.1101/2025.05.05.652253

**Authors:** Cameron J. Douglas, Ciaran P. Seath

**Affiliations:** Department of Chemistry, Wertheim UF Scripps, Jupiter, Florida, 33418, United States; The Skaggs Graduate School of Chemical and Biological Sciences, 120 Scripps Way, Jupiter, FL 33458, USA

## Abstract

Omics analysis has become an indispensable tool for researchers in the life sciences, enabling the study of DNA, RNA, and proteins and how they respond to cellular stimulus. Many methods of data analysis exist for the generation and characterization of gene lists, however, selection of genes for further investigation is still heavily influenced by prior knowledge, with practitioners often studying well characterized genes, reinforcing bias in the literature. Here, we have developed an open-source, R package for impartial ranking of gene lists derived from omics analysis that we term Deciphering Scientific Discoveries (DeSciDe). We applied a pipeline that sorts a gene list first by precedence, which we define as co-occurrence of the gene with pre-defined search terms in publications. We then rank gene lists by connectivity, an underutilized metric for how related a gene is to other enriched genes. The combination of these rankings by scatterplot provides a method for gene selection by simple visual analysis. We apply this analysis methods to published Omics datasets, identifying novel avenues for investigation. Further, using this method we have been able to assign a novel loss of function role for the histone mutation H2A E92K.

## INTRODUCTION

The study of biochemical processes is increasingly performed using large scale omics analyses to survey changes in RNA or protein expression (RNA-seq/proteomics). These experiments generate large data sets that require robust filtering and analysis to determine broader biological implications. A suite of effective tools has been developed to this effect such as STRING, a database of known and predicted protein-protein interactions, gene ontology (GO)^1,2^, a knowledge base about the functions of genes, and gene set enrichment analysis (GSEA)^3^, a computational method that determines whether an a priori defined set of genes shows statistically significant, concordant differences between two biological states (Figure 1A)^2–6^. These open-source tools have become an essential resource and are widely used throughout the community. These tools are well suited to uncovering relationships between enriched genes but are not able to study relationships between enriched genes and the cellular stimulus under study, leading to a degree of manual curation that results in human bias towards the study of well characterised genes. This bias perpetuates the continued study of a subset of genes, leaving many more underexplored. When deploying more complex omics experiments, such as proximity proteomics, where nearby proteins to a protein of interest are identified^7^ or CRISPR screens, where ranked lists of genes that confer sensitivity or resistance to a biological challenge of interest are identified from cells with genetically encoded perturbations^8^, the importance of data curation is even more critical as false positives are more common.

**Figure 1.**
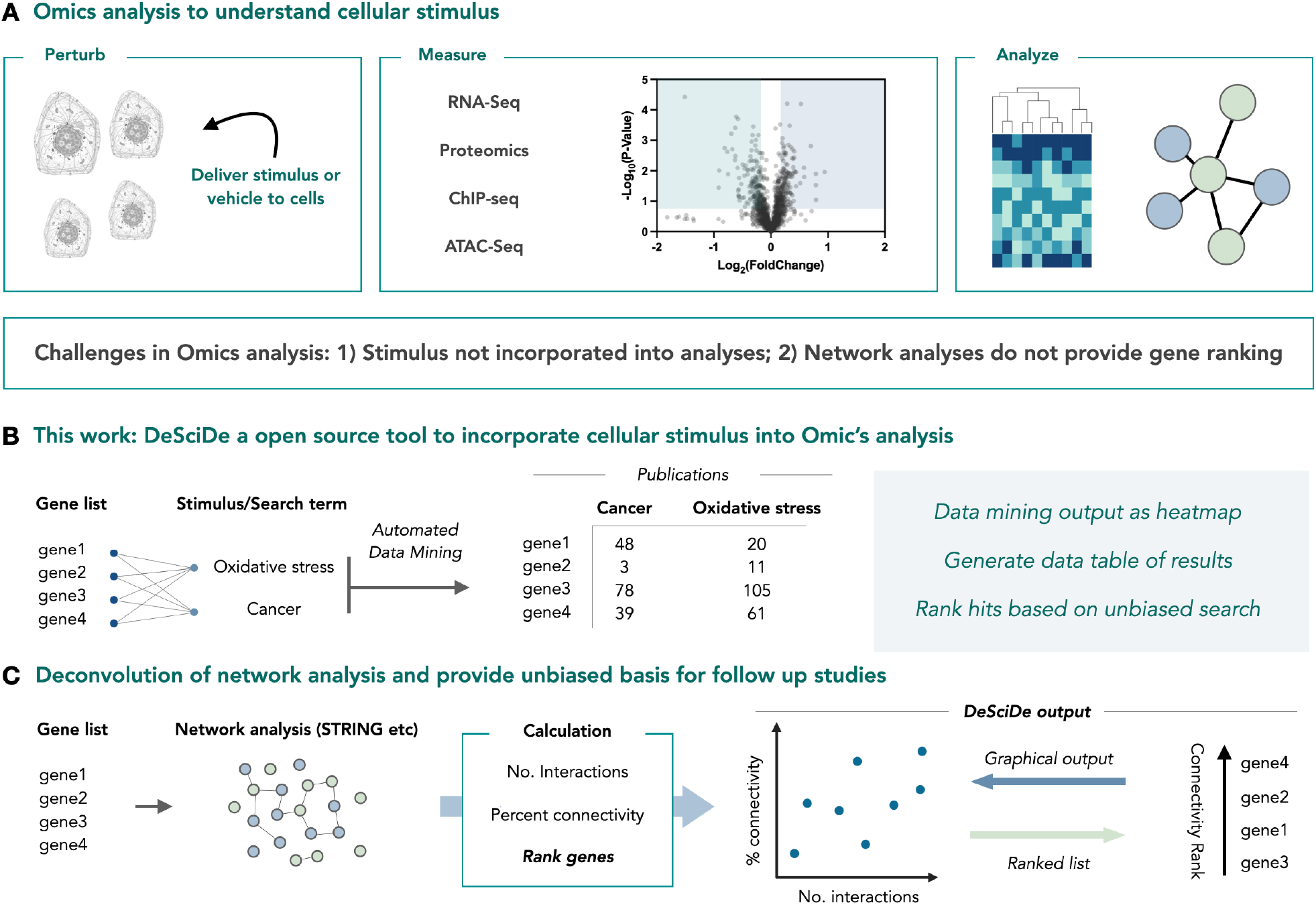
An open-source pipeline for unbiased omics analysis. **a**, Omics analysis is often used to understand how stimulus can change cellular biology. **b**, DeSciDe performs automated literature data mining to rank gene lists for precedence within a particular field or in relation to a particular stimulus or disease. **c**, DeSciDe ranks genes according to their STRING connectivity, providing a ranked list of genes. Connectivity analysis can provide a numerical basis for choosing genes for follow up studies.

When studying gene lists from omics analyses, it can be valuable to search for interactions between enriched genes to aid users in assigning molecular mechanisms. These physical interactions are typically explored using STRING analysis, where genes are plotted as interconnected nodes. This tool is enabling when studying small lists of genes, but the graphical output becomes unwieldy and challenging to deconvolute when large numbers of interconnected genes are present, limiting the use of the tool when analysing lists with greater than 50 hits.

Based on these limitations, end users of omics methods have an urgent need for tools to aid in unbiased selection of gene hits for follow up studies. In considering this need we identified several requirements; 1) a method must incorporate the cellular stimulus or pathway that is being studied; 2) the method should be able to assign value to interactions between genes; 3) the program must be readily available and applicable to many different experimental types.

To address these requirements, we have developed an R package to enable the analysis of gene lists that are experimentally associated with a relevant search term. Our package, Deciphering Scientific Discoveries (DeSciDe), can incorporate any number of cellular stimuli and cross-reference them against lists of enriched genes. Based on the co-occurrence of the gene and the cellular stimulus, a hit is assigned as either strongly or poorly associated with a particular search term. Further, our system computes network connectivity metrics from the gene interaction networks provided by the STRING database, producing a quantitative evaluation of connectivity. The gene lists can then be sorted based on two key values: interconnectivity and precedence. Through analysis of existing datasets, we demonstrate that connectivity is a viable metric for selecting genes for follow up investigation, and when combined with the quantification of literature precedence, can provide a systematic and objective method for gene selection. We anticipate that such a tool will be valuable for myriad applications across the biological sciences.

## RESULTS AND DISCUSSION

We began developing a tool to search gene lists against any desired term that could represent a cellular stimulus (Figure 1B). This was achieved by cross searching every gene in the given list against each search term using PubMed abstracts as the database. The resulting references are reported in a table as the number of references per gene and search term combination. Additionally, we incorporate a graphical representation of the citations in the form of a heatmap. The heatmap shows number of references for each stimulus/search term, which can be used to visualize the top “hits” based on literature precedence for multiple search terms at once. DeSciDe then ranks the genes based on literature precedence. This can be done using two different methods: weighted and total. The default method is weighted, in which the gene list is filtered by the number of publications associated with the first input term followed by filtering for the subsequent terms in the order provided. This method allows the user to prioritize highly specific search terms (e.g., histone H3 K23 acetylation), while still incorporating broader cellular contexts (e.g., cancer) in their terms list without biasing the results toward the term with the highest numbers of publications. Alternatively, ranking by total number of publications across all search terms for a gene can be conducted when users do not deem it necessary to prioritize a specific cellular stimulus context.

Next, we sought to establish a metric for quantifying and ranking gene interconnectivity, which we suggest may be valuable and generally applicable for hit selection following omic analyses. We incorporated existing STRING interaction networks and quantified each gene according to two criteria: number of interactions (known as *degree* in graph theory), and connectivity (known as *clustering coefficient* in graph theory) (Figure 1C). The number of interactions is straightforward, representing the number of connections made by each node. A gene’s connectivity score represents the percentage of existing connections compared to the theoretical maximum number of connections in the subnetwork spanned by the respective node (the network comprised of the node and its neighbours). We then compile the gene lists and their network properties in a table that is filterable. By default, the genes are ranked first by the number of connections and then by percent connectivity. The package also produces a scatterplot showing the number of interactions versus the percent connectivity. This type of plot provides a visual representation of how interconnected the genes in the list are. Genes in the bottom left of the plot have few connections and low connectivity, whereas the top right corner contains the most connected set of genes (Figure 1C).

The final component of the application is the combination of precedence and connectivity rankings. Here, we create a scatterplot that displays the rank order of precedence versus the rank order of connectivity. Since rank order arranges values from high to low, in this visualization, hits that have high precedence and high connectivity appear close to the origin. By default, DeSciDe classifies these genes based on a 20% threshold of total genes in the list with high-connectivity, high-precedence genes falling in the top 20^th^ percentile of ranked genes in both lists and high-connectivity, low-precedence genes falling in the top 20^th^ percentile of connectivity and the bottom 20^th^ percentile in precedence. This threshold can be adjusted by the user as deemed fit for their analysis. We found this to be a broadly useful visualization for hit selection. To illustrate how DeSciDe might be used we applied our workflow to four case studies.

The plots produced by the DeSciDe analysis pipeline can be easily exported and saved for use in presentations and publications. Examples of the plots produced by default running of DeSciDe can be seen in supporting information (Figures S1-S5). Additionally, the data tables of results can be saved as TSV, CSV, or Excel files for further analysis or for generating revised figures.

We began by analysing proximity proteomics data sets. We chose these as examples as they are uniquely suited to this analysis for several reasons: 1) the bait (or protein of interest bearing the proximity labelling enzyme or catalyst) is not typically included in standard analysis as it is often spatial^9^ or abiological (i.e., a small molecule^10^), and 2) these experiments are typically designed to look for unknown interactions. Therefore, careful filtering of gene lists is required before choosing genes of interest for further investigation, adding an element of human bias.

First, we analysed a proximity proteomics dataset published by Geri et al. that describes the interactome of the receptor PDL1 on Jurkat cells^9^. PDL1 plays an important role in the immune system as a “save me” signal, stopping immune cells from attacking healthy cells. As PDL1 is frequently expressed by cancer cells to the immune system, it has become an active oncology target^11^. The purpose of this experiment was to identify novel interactors of PDL1 that may play a role in immune oncology. From the gene list provided, we filtered for all differentially enriched genes that met statistical significance (41 total) and passed them through the DeSciDe pipeline (Figure 2A) with the keyword’s *cancer, immunology*, and *checkpoint blockade*. STRING analysis of the 41 hits was informative, with 12 genes in the list having no known interactions and 24 genes having at least three known interactors within the data set (Figure 2B). To illustrate how connectivity is calculated from STRING data, nodes surrounding FCER2 and HLA-B are shown in Figure 2C. DeSciDe ranks these genes by connectivity (Figure 2D), placing ICAM1, CD40, and CD274(PDL1) as the top three hits. Of these three, CD274 is the bait protein, and CD40 and ICAM1 have both been validated to colocalize with PDL1 in immune synapses and are themselves targets for immune based therapies^13–15^.

**Figure 2.**
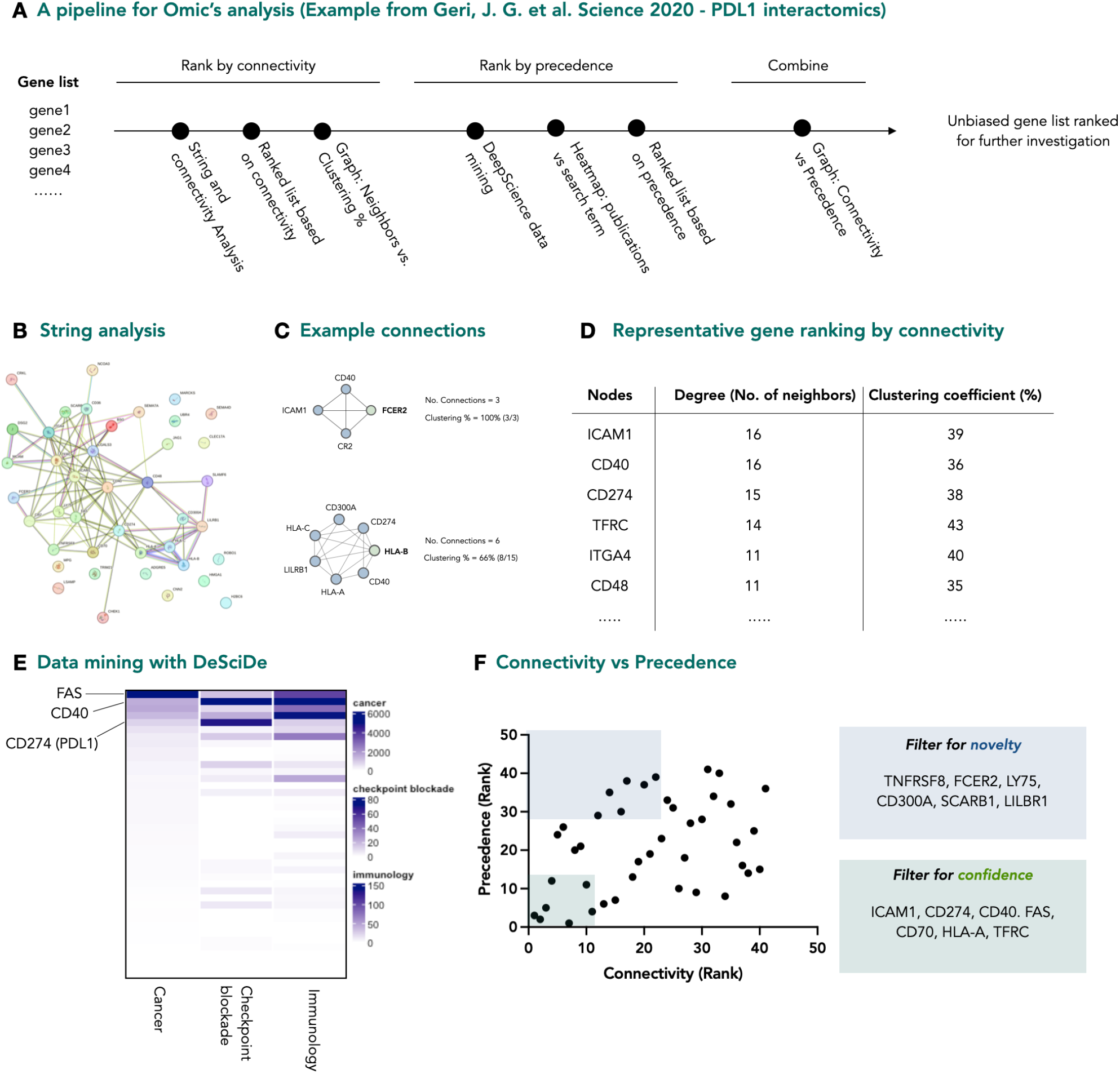
Pipeline for analysis of a proximity proteomics dataset. **a**, Graphical representation of DeSciDe pipeline for unbiased ranking of gene lists. **b**, STRING analysis of PDL1 interactomics data set, published by Geri et al.^9^ **c**, Example of how connectivity analysis is performed. **d**, PDL1 interactome ranked by connectivity (top 6 genes shown). **e**, Heatmap showing results of DeSciDe data mining against the search terms “cancer” “immunology” and “checkpoint blockade”. **f**, Scatterplot of genes ranked for connectivity vs precedence with suggested alternate genes for investigation highlighted in boxes. Graphs made in Prism from exported DeSciDe data.

Next, using DeSciDe datamining, we cross referenced the gene list with the search terms described above and plotted via heatmap (Figure 2E). These data clearly show that the genes within this data set have significant overlap with the search terms *immunology* and *cancer*, but fewer with the more specific term *checkpoint blockade*. Our application makes it trivial to identify related genes from the heatmap. The ranked lists were then combined and displayed as a scatterplot of precedence vs. connectivity (Figure 2F). The genes of highest connectivity and highest precedence are located near the origin. In this data set, that includes the previously discussed ICAM1, CD40, and PDL1, in addition to CD70, HLA-A, and the death receptor FAS. Based on previous reports in the area and the precedence from the data mining, all the genes within this sector are confident hits^16–19^. With this knowledge, it can be assumed that connectivity is a reasonable metric to rank genes. If this is correct, then moving to the top left quadrant, where connectivity remains high, but the genes have far fewer precedented reports relating to the three search terms, may provide novel targets for investigation. In this area, we found 7 genes (TNFRSF8, FCER2, LY75, CD300A, SCARB1, LILBR1) that are candidates to be novel interactors, with less known about their involvement in PDL1 based checkpoint blockade.

In our next case study, we examined a proximity proteomics dataset with a significantly more complex interactome. The experiment, published recently by the laboratories of MacMillan and Muir, described how a somatic mutation on histone H2A disrupts the nucleosome microenvironment (Figure 3A)^12^. This type of hypothesis generating experiment is a good match for our data analysis pipeline as the proteomics data can lead to several areas of study and is easily influenced by inherent bias. In the original study the authors identified several enriched genes (SIRT6, DNMT3A/B, BRD2/3/4) for further investigation. When analysing this data set with our pipeline, we found that these genes were among the most well studied (by searching the terms *chromatin, nucleosome*, and *acidic patch*) ranking 2^nd^, 3^rd^, 4^th^, 11^th^, and 12^th^ of all genes in our measure of precedence (Figure 3B). Visual inspection of STRING analysis for this data set was not instructive due to the density of the interaction networks (Figure 3C). However, connectivity analysis suggested a set of genes that were not implicated in the original study (Figure 3D & 3E).

**Figure 3.**
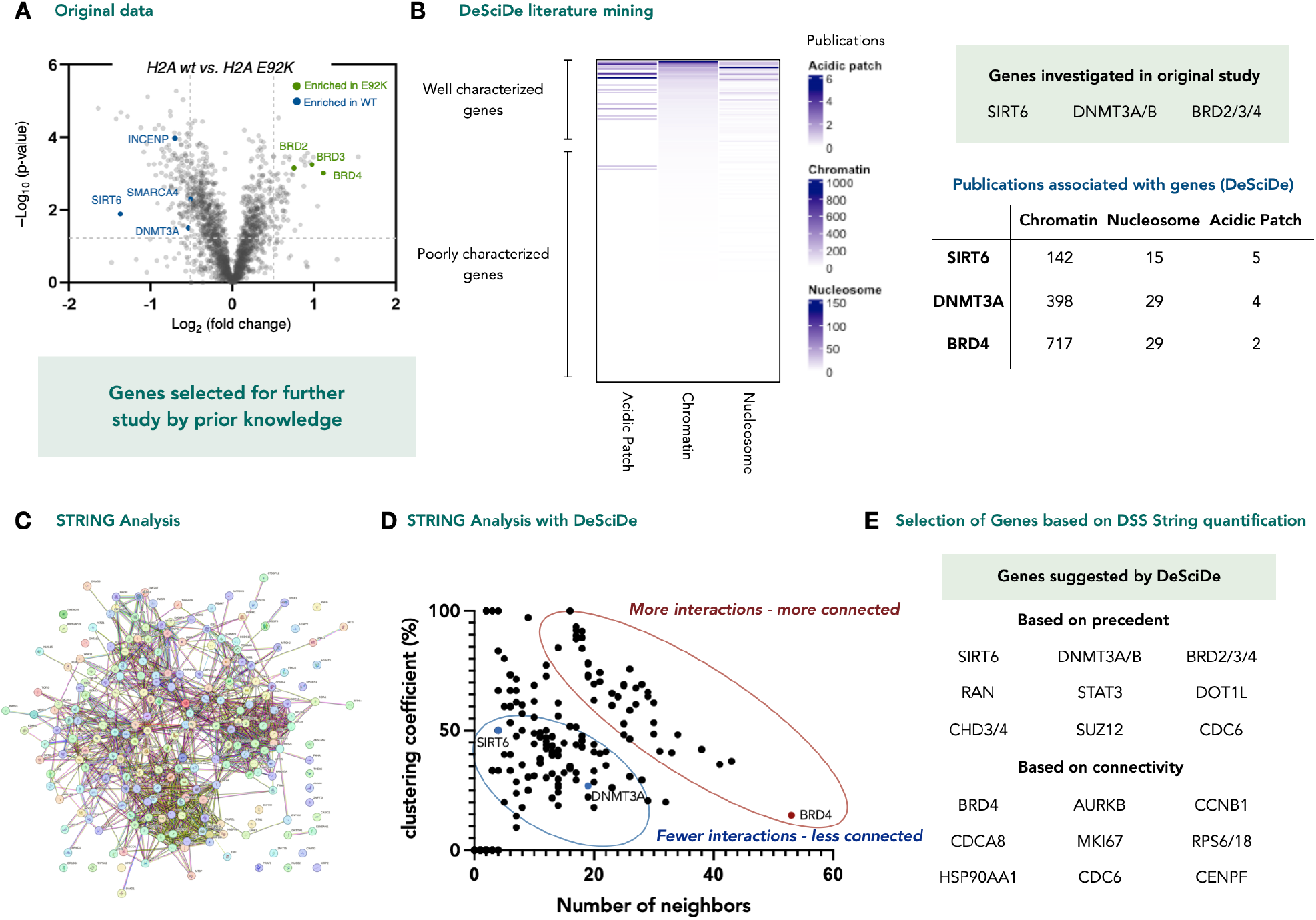
Re-analysis of proximity proteomics data studying the effect of somatic mutations on the nucleosome acidic patch. **a**, Original published dataset. Genes highlighted were validated or taken for further investigation^12^. **b**, Heatmap derived from DeSciDe analysis using the search terms “chromatin” “nucleosome” and “acidic patch”. The genes taken on for further study were in the top 5 most studied genes (in the context of the three search terms employed). **c**, STRING analysis of significantly enriched genes is too complex for visual interpretation. **d**, Graphical interpretation of STRING analysis via DeSciDe computed connectivity. Genes are more readily visualized as being highly connected. **e**, Unbiased gene selection by DeSciDe, sorting for either precedent or connectivity. **f**, DeSciDe plotting of connectivity vs precedence provides new avenues of investigation. **g**, DeSciDe analysis suggests genes related to regulation of cell cycle for further analysis, a novel phenotype for acidic patch mutations. Graphs made in Prism from exported DeSciDe data.

Plotting the ranked lists against each other proved to be enlightening, with both critical quadrants i.e., nearest the origin (high confidence genes) and the top left quadrant (highly connected but less well studied in this context), both suggesting investigation of genes relating to regulation of the cell cycle (Figure 4A&B).

**Figure 4.**
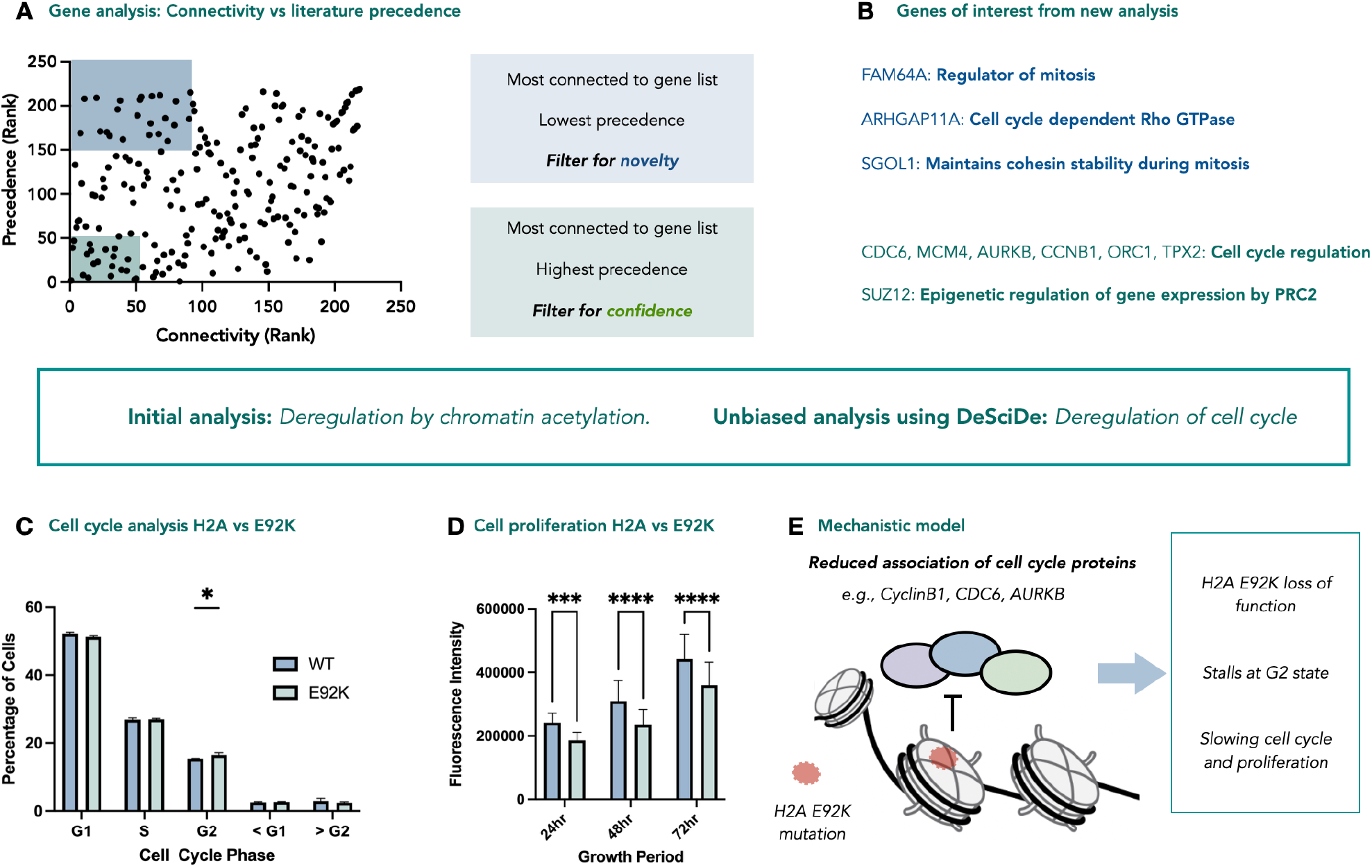
Re-analysis of proximity proteomics data studying the effect of somatic mutations on the nucleosome acidic patch. **a**, DeSciDe plotting of connectivity vs precedence provides new avenues of investigation. **b**, DeSciDe analysis suggests genes related to regulation of cell cycle for further analysis, a novel phenotype for acidic patch mutations. Graphs made in Prism from exported DeSciDe data. **c**, Cell-cycle analysis via flow cytometry using propidium iodide (n=3, 50,000 cells counted per replicate, whiskers represent standard deviation). P-value for 1.1% difference in G2 phase = 0.0221. **d**, Cell proliferation data over 72h comparing HEK293T expressing H2A or H2AE92K plated at 10,000 cells/well in a 96-well plate at time = 0hr (n=30, whiskers represent standard deviation, data graphed represents mean fluorescent intensity at indicated timepoint). **e**, Mechanistic model for cell-cycle stalling and reduced proliferation for E92K mutation. *P < 0.05, *** P < 0.001, **** P < 0.0001.

Recently, McGinty and co-workers demonstrated the role of the acidic patch in coordinating VRK1 phosphorylation of H3T3 during cell division^20^. Furthermore, pathogenic mutations on VRK1 were shown to disrupt this interaction, providing a molecular basis for how these rare mutants may cause rare adult-onset distal spinal muscular atrophy. Our reanalysed data also suggest that the acidic patch may play a role in cellular division, and that the E92K mutation may lead to deregulation of this critical cellular pathway.

We further investigated this by performing cell-cycle analysis using propidium iodide in HEK293T cells stably expressing H2A or H2A E92K. We observed the mutant cell line contains a higher concentration of cells in the G2 phase, suggesting that the small proportion of mutated nucleosomes confer a subtle change in the cell cycle (Figure 4C). Based on this stalling, we anticipated that the mutant cell line would show reduced proliferation compared to the wild-type cell line. Cell proliferation assays of both cell lines over 72 hours showed the E92K mutation significantly decreased proliferation, in line with our hypothesis (Figure 4D). Based on these data, we can assign a new role for the E92K acidic patch mutation. The previously reported chemoproteomics data shows that the mutation disrupts interactions between the nucleosome acidic patch and cell cycle proteins AURKB, CDC6, and CyclinB1, which all play a significant role in the G2-M transition during cell division^21–23^. This disruption leads to stalling in the G2 phase of cell division and subsequent reduction in proliferation (Figure 4E). The data suggests these effects are subtle, likely due to the relatively low incorporation (approx. 1 in 16 nucleosomes), and that it is likely that only one face of each mutant nucleosome bears the mutation. However, this effect is in line with the chemoproteomics and ATAC-seq data previously reported^12^. This example indicates that unbiased analysis using a program such as DeSciDe can provide different avenues of investigation that may be overlooked in favour of genes that are highly represented in the literature.

Furthermore, we demonstrate that DeSciDe is widely applicable to omics analysis by reanalysing publicly available datasets from RNA-seq, global proteomics, CRISPR screens, and ATAC-seq experiments. A recent RNA-seq experiment published by Ma et. al., was deployed to study differential splicing in the context of TDP-43 deficient FTD-ALS^24^. The authors chose the mRNA UNC13A for follow up studies, demonstrating its role in ALS pathology (Figure 5A). Plotting connectivity vs precedence using the search terms RNA-splicing, ALS, and TDP-43 suggested UNC13A as the highest confidence hit (most connected and most precedented), illustrating that this bioinformatics methodology can recapitulate complex data analyses without prior knowledge of the field (Figure 5B). Further, based on the scatterplot, we can suggest alternative genes for investigation that either cluster around UNC13A or have less precedented associations with the search terms, which may not be obvious candidates for follow-up studies (Figure 5C).

**Figure 5.**
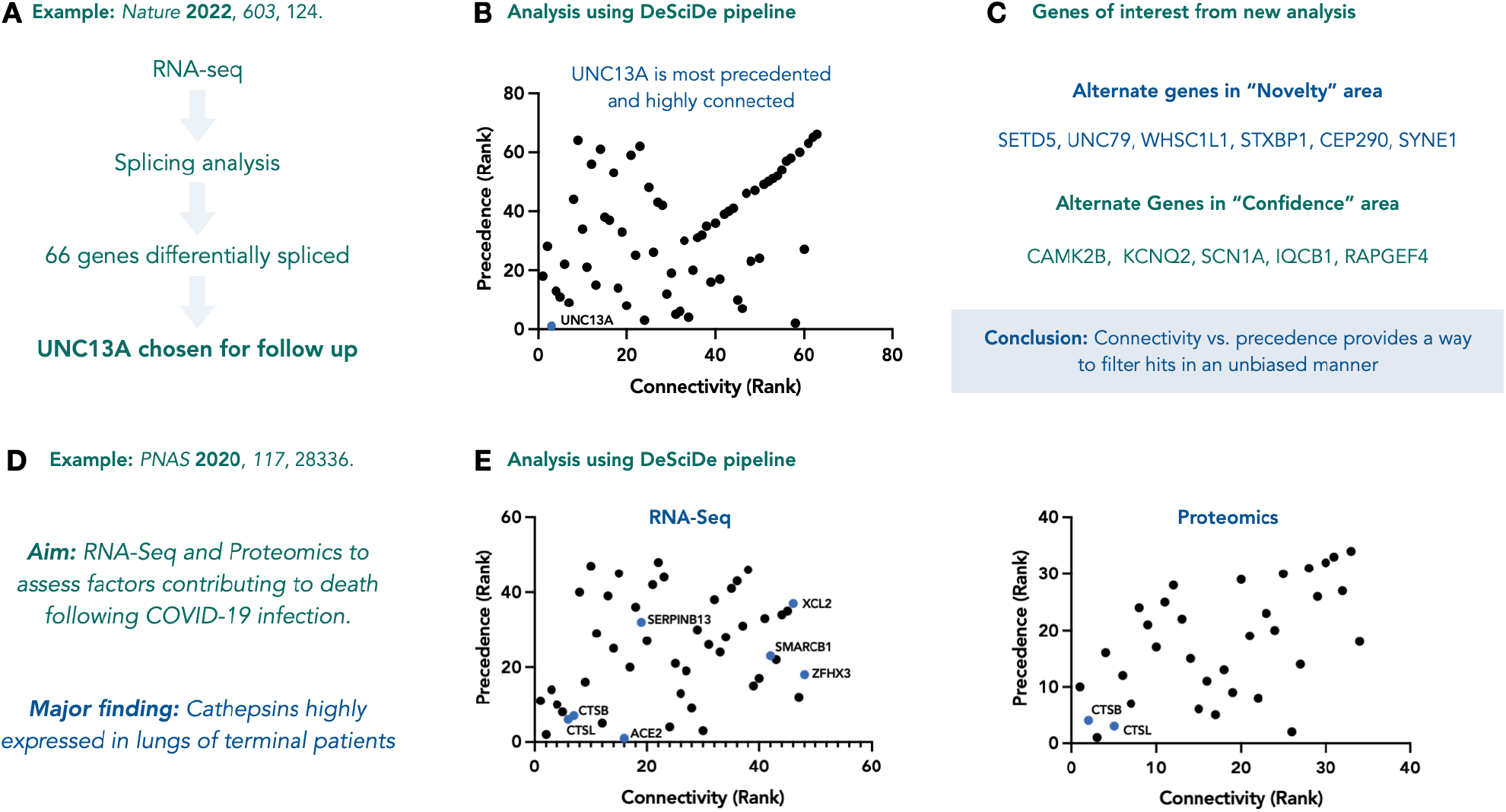
Analysis of RNA-seq data using DeSciDe. **a**, Brief outline of RNA-seq experiment performed in the study by Ma et al.^24^ **b**, DeSciDe analysis suggests UNC13A as the highest confidence hit, recapitulating the authors analysis in an unbiased manner. **c**, DeSciDe analysis can suggest alternate genes for further analysis based on novelty or confidence. **d**, Aim of proteomics and transcriptomics experiments re-examined using DeSciDe. **e**, Both proteomics and transcriptomics analyses via DeSciDe arrive at the same conclusion as the authors. Many high confidence genes remain unexplored within both studies^25^. Graphs made in Prism from exported DeSciDe data.

Finally, we analysed a 2020 study on the proteomic and transcriptomic host response to COVID-19^25^. One key aim from these experiments was to identify factors that contribute to fatality following infection (Figure 5D). From 4065 and 637 differentially expressed genes across RNA-seq and proteomic datasets, respectively, the authors highlighted expression of cathepsins as a marker for poor prognosis. DeSciDe analysis also points towards CTSB and CTSL as high confidence hits in the quadrant closest to the origin in both RNA-seq and proteomics datasets (Figure 5E). Once again, these data suggest that analysis through the DeSciDe pipeline and using connectivity and literature datamining to rank hits is a viable and useful bioinformatics method for unbiased analysis of gene lists.

While this method of data analysis appears to be powerful for ranking genes in an unbiased way, some deficiencies remain. Connectivity is based upon and intrinsically related to precedence, where more connections have been reported for more “popular” genes. So completely uncharacterized genes will still be ignored using this analysis. Further, the search function within this application cannot filter for the most relevant journal articles, only those that contain the gene name and the manually selected search terms, so some articles may not actually show a meaningful connection. Finally, these analyses do not include enrichment fold change, which is often a metric used for gene selection. We are currently working on methods to improve the analysis pipeline to solve some of these issues.

## CONCLUSION

In summary, we have developed an open-source R package for unbiased analysis of gene sets from omics experiments. The application can rank genes by how connected they are to the rest of the dataset and by how well they are associated in the literature with predefined search terms. The combination of these two rankings provides a valuable scatterplot that can be used to identify high confidence genes for further investigation. Using this method, we reanalyse a proximity proteomics data set and identify new biological implications of H2A E92K mutations. We believe this application will find broad usage within the life sciences and will aid researchers in identifying new avenues for biological investigation. The code of the application is freely available under the MIT *License* at https://github.com/camdouglas/DeSciDe.

## ASSOCIATED CONTENT

## Supporting Information

The Supporting Information contains a vignette for detailed instruction on how to use the DeSciDe package and tables of the genes selected from select publications and their DeSciDe results (XLXS). DeSciDe is available for download on CRAN at https://cran.r-project.org/web/packages/DeSciDe/index.html. Code can be accessed at https://github.com/camdouglas/DeSciDe. The Supporting Information is available free of charge on the ACS Publications website.

## AUTHOR INFORMATION

## Authors

Cameron Douglas – Department of Chemistry, Wertheim UF Scripps Institute, Jupiter, Florida, 33418, United States. The Skaggs Graduate School of Chemical and Biological Sciences, 120 Scripps Way, Jupiter, FL 33458, USA

## ACKNOWLEDGMENT

CPS would like to thank Wertheim UF Scripps for startup funding. This work was supported by NIH (1R35GM150765-01 to CPS)

