## Supplementary Figures and data for "DeSciDe: A tool for unbiased literature searching and gene list curation unveils a new role for the acidic patch mutation H2A E92K"

#### *Methods*

##### Flow Cytometry Cell Cycle Analysis

HEK293T cells expressing either H2A or H2A E92K were seeded at 5 x 106 cells per 10cm plate. At 24hrs cells were collected and fixed with 70% -20C Ethanol. Fixed cells were pelleted and resuspended in FxCycle PI/RNase Staining Solution (Invitrogen) and incubated for 30min at room temperature. Cells were analysed by flow cytometry on a BD LSRII flow cytometer with the yellow-green laser and 50,000 events were recorded for each replicate (n=3). Cell cycle phases were gated and quantified in FlowJo. Statistical analysis conducted in Prism GraphPad by a 2-way ANOVA.

##### Cell Proliferation Assay

HEK293T cells expressing either H2A or H2A E92K were seeded at 10,000 cells/well in 30 wells of a 96-well clear bottom plate with Fluorobrite DMEM (Gibco) supplemented with 10% FBS. At 24hrs, 48hrs, and 72hrs post seeding, the plates were read out by the addition of 2x CellTiter-Fluor Cell Viability Assay substrate (Promega). Cells were incubated for 30min at 37C and then fluorescence (390/505nm) was measured 3 times for each plate on a plate reader. Fluorescence intensity for each replicate was averaged, background signal from empty wells was subtracted, and results were plotted in Prism GraphPad and P-values determined by a 2-way ANOVA.

#### *Supplementary Figures*

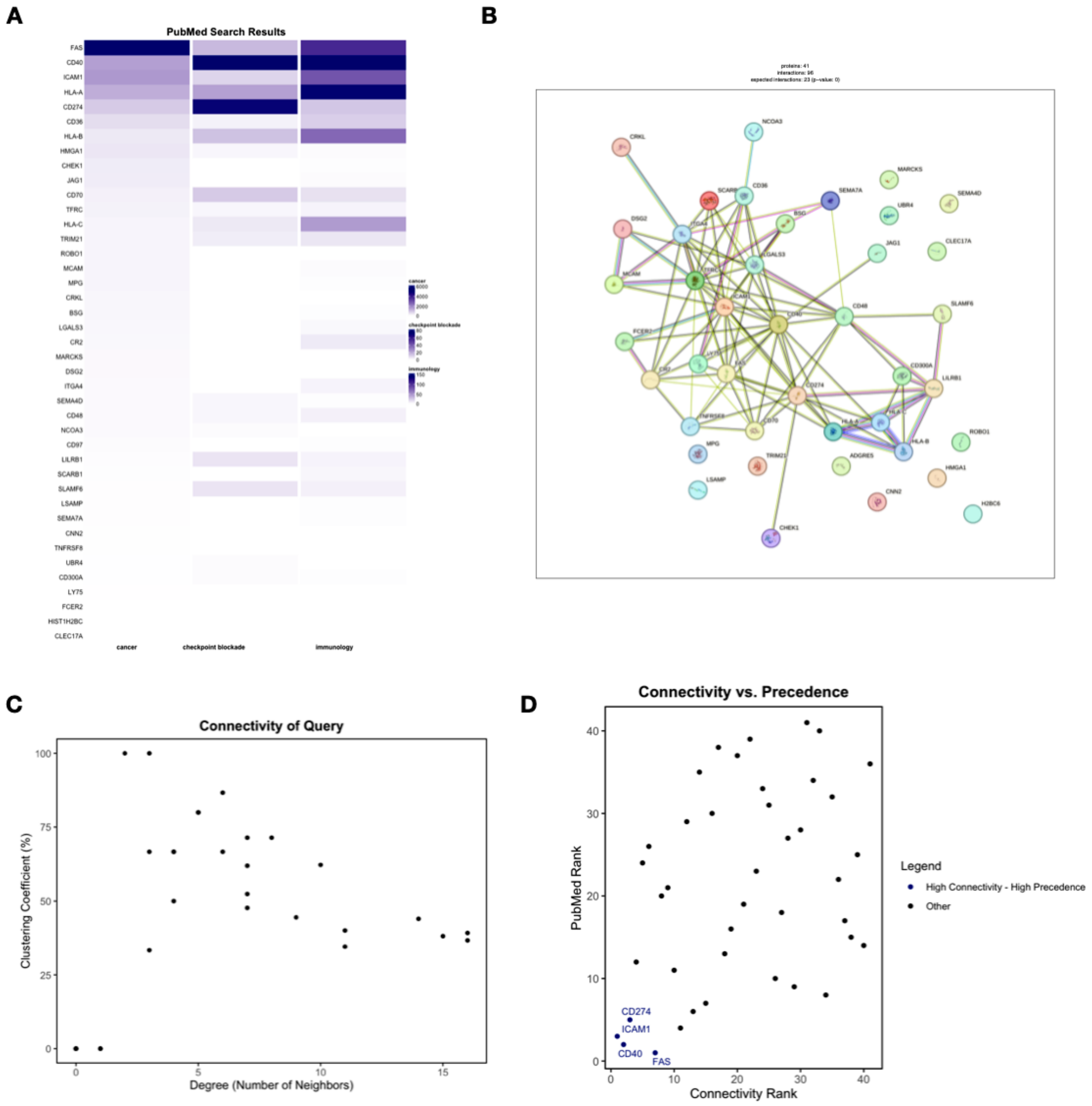

**Figure S1.** Plots generated by DeSciDe for Geri et al. 2020 data.

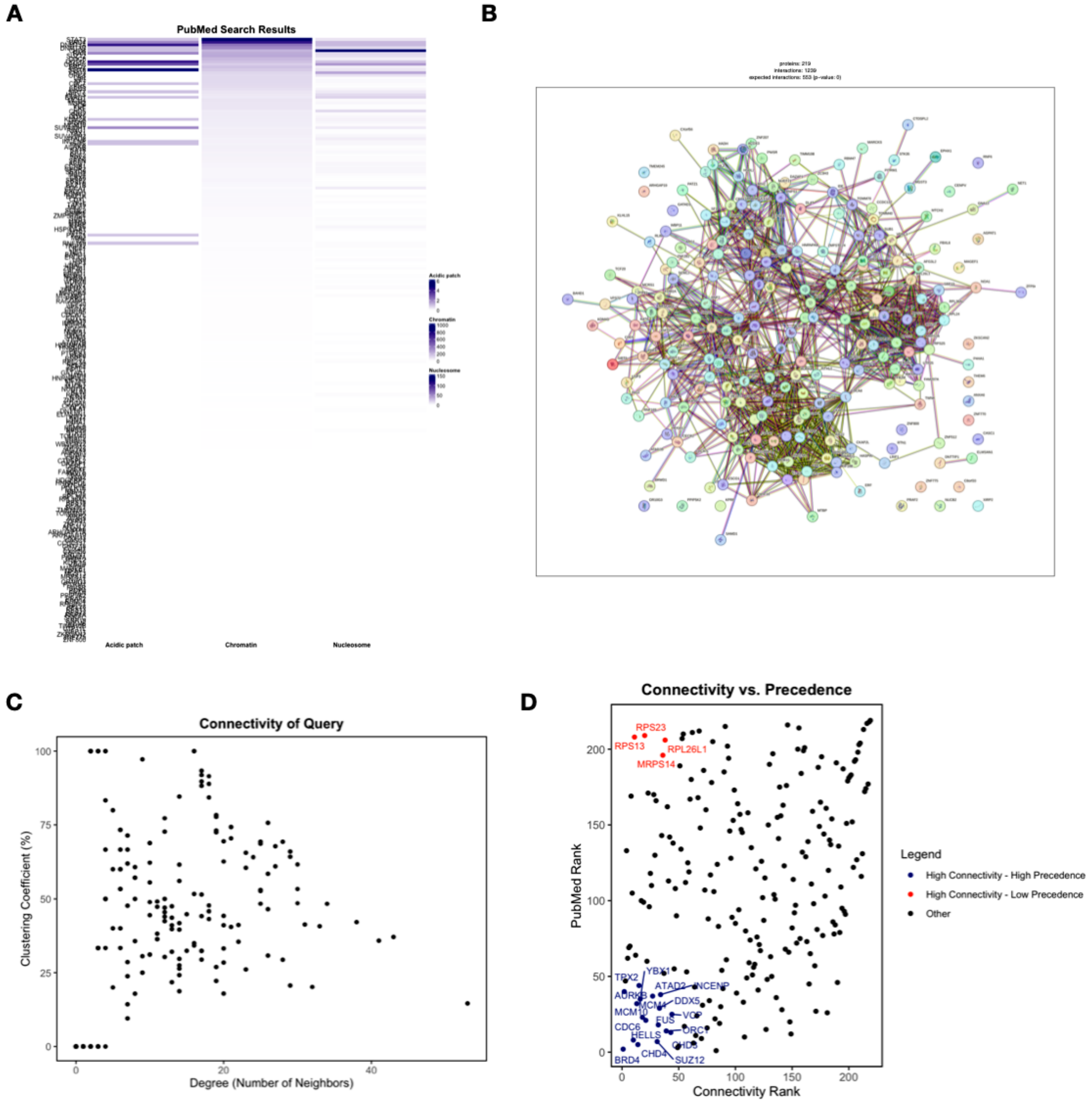

**Figure S2.** Plots generated by DeSciDe for Seath et al. 2023 data.

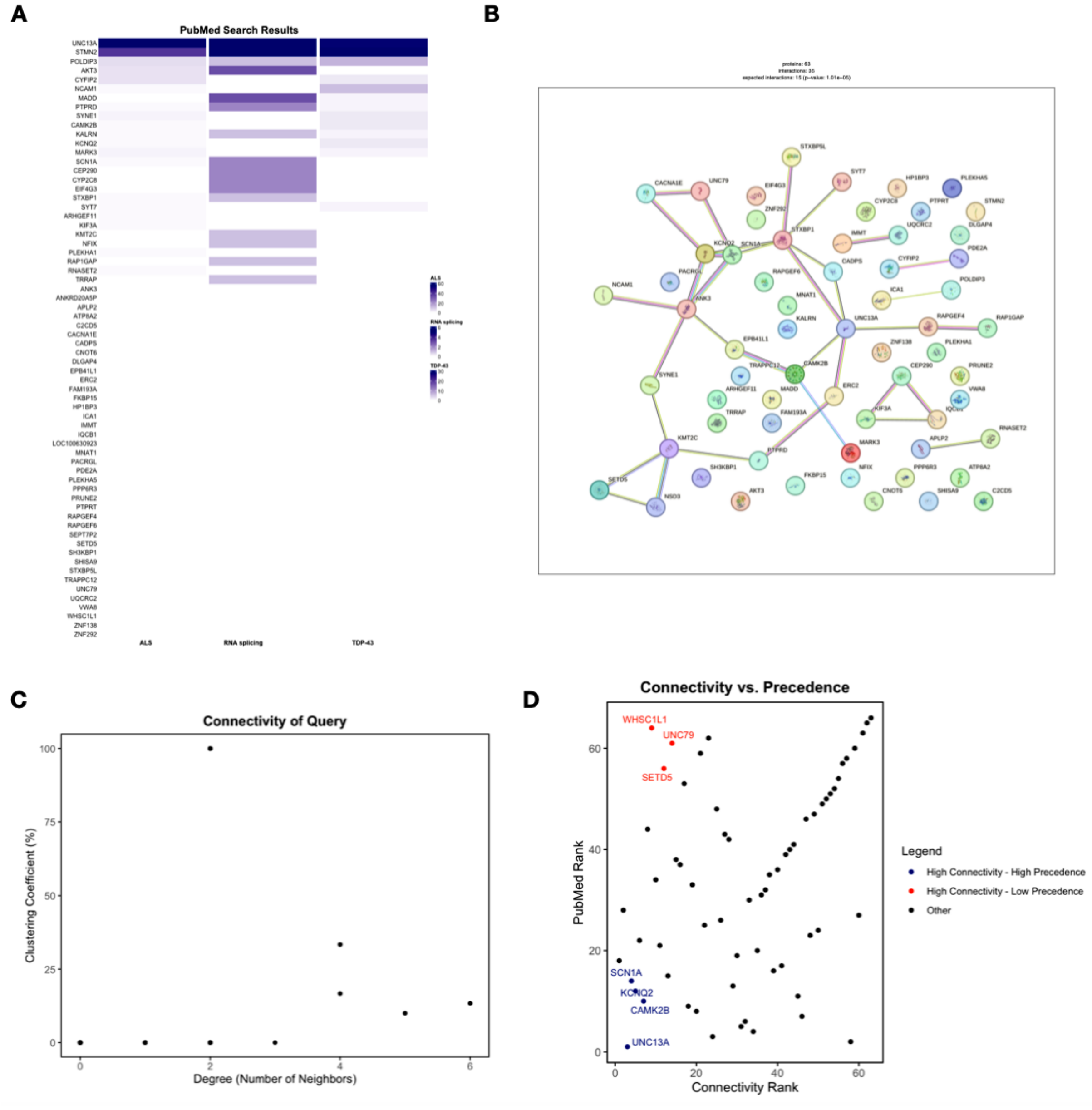

**Figure S3.** Plots generated by DeSciDe for Ma et al. 2022 data.

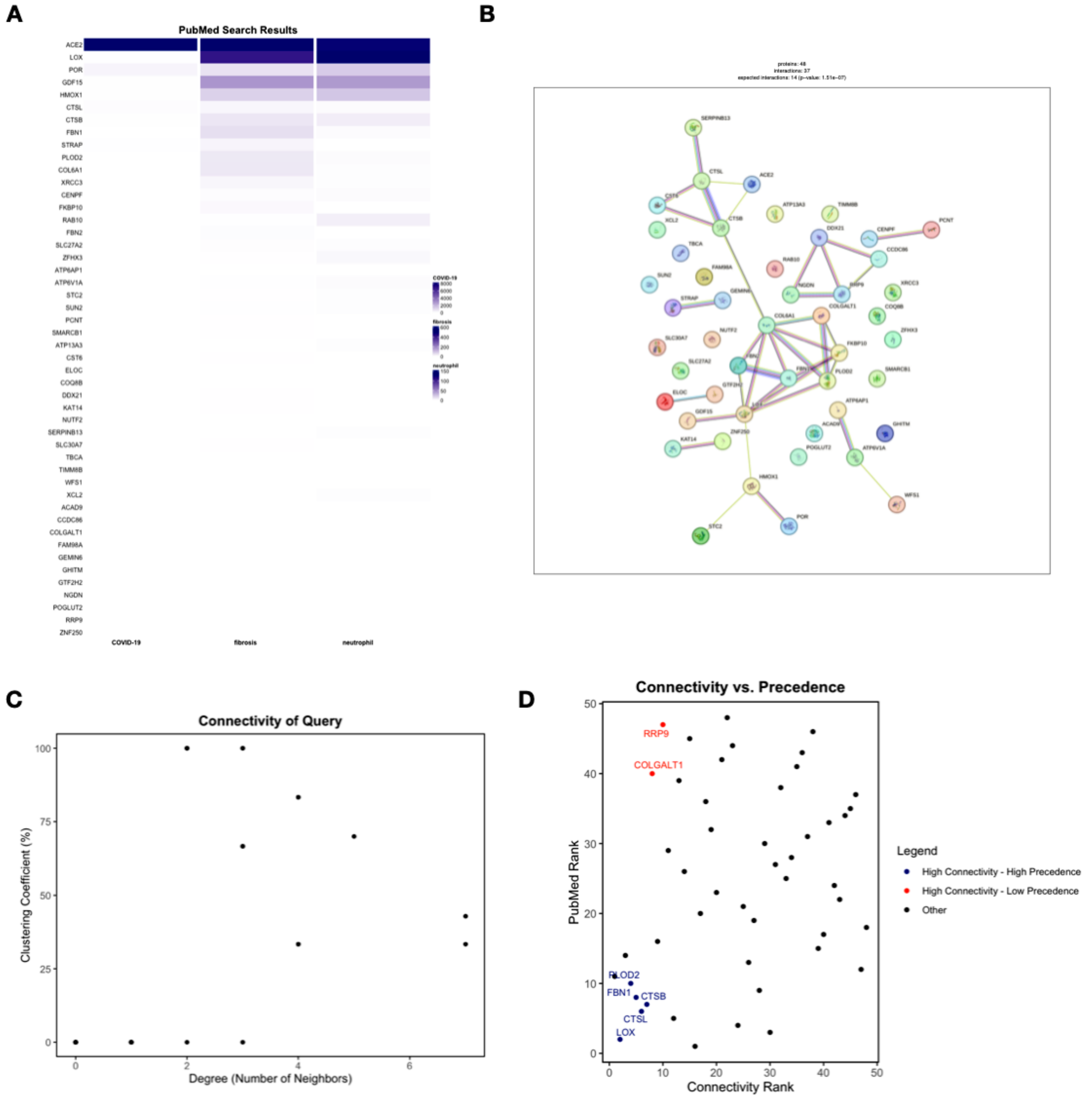

**Figure S4.** Plots generated by DeSciDe for Wu et al. 2020 RNA-seq data.

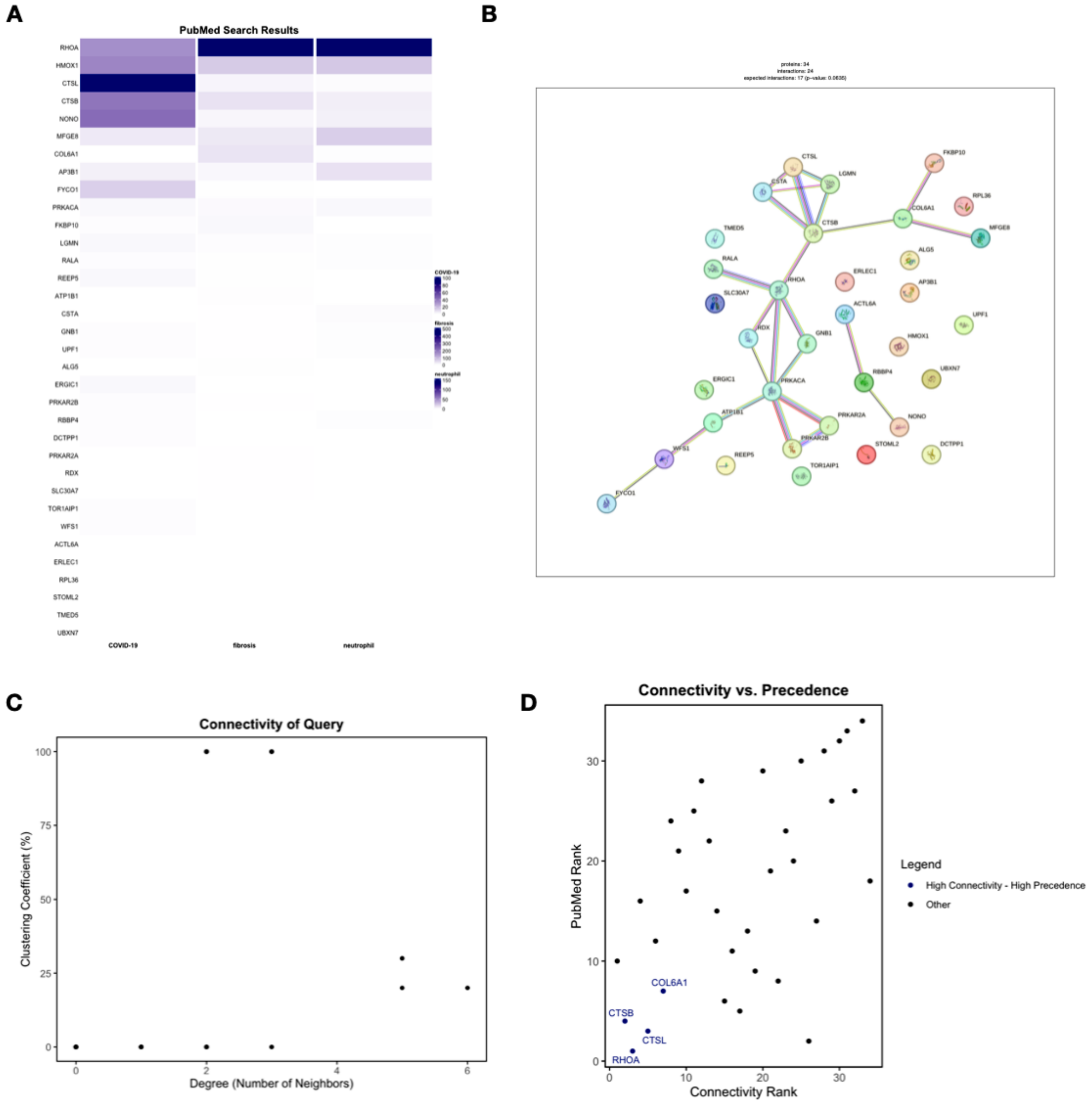

**Figure S5.** Plots generated by DeSciDe for Wu et al. 2020 proteomics data.

### DeSciDe Vignette

#### DeSciDe (Deciphering Scientific Discoveries)

DeSciDe is a package designed to help streamline omics data analysis. Many methods of data analysis exist for the generation and characterization of gene lists, however, selection of genes for further investigation is still heavily influenced by prior knowledge, with practitioners often studying well characterized genes, reinforcing bias in the literature. This package aims to aid in the identification of both well-studied, high-confidence hits as well as novel hits that may be overlooked due to lack of prior literature precedence.

This package takes a curated list of genes from a user's omics dataset and a list of cellular stimuli or cellular contexts pertaining to the experiment at hand. The list of genes is searched in the STRING database, and informative metrics are calculated and used to rank the gene list by network connectivity. Then the genes list and terms list are searched for co-occurrence of each gene and search term combination to identify the literature precedence of each gene in the context of the search terms provided. The PubMed results are then used to rank the genes list by number of publications associated with the search terms.

The two ranks for each gene are then plotted on a scatter plot to visualize the relationship of the genes' literature precedence and network connectivity. This visual aid can be used to identify highly connected, well-studied genes that serve as high confidence hits clustered around the origin and highly connected, low precedence genes that serve as novel hits clustered in the top left of the graph. The highly connected, low precedence genes are known to interact with the other genes in the list, but have not been studied in the same experimental context, providing novel targets to pursue for follow up studies.

Additional graphical outputs are generated to visualize the STRING network and PubMed results. The package has also been set up to allow the user to change various steps of the analysis to fit their needs, such as searching STRING for all connections or just physical, adjusting the threshold of classification of genes, exporting figures and data tables for use in publications. Below, we highlight how to implement DeSciDe to study an example data set.

##### Example Usage of DeSciDe

For the following examples we will use a list of 40 genes and 3 search terms. We will call these lists "genes" and "terms". Here we import the lists from CSV files, however the user can choose to manually create these lists or import how they see best fit.

```
# Import genes list and terms list from CSV
genes <- read.csv("genes.csv", header = FALSE)[[1]]
terms <- read.csv("terms.csv", header = FALSE)[[1]]
#> Warning in read.table(file = file, header = header, sep = sep, quote = quote, :
#> incomplete final line found by readTableHeader on 'terms.csv'
```

We now have a list of our genes and terms to execute in our code:

```
genes
#>  [1] "BRD4"     "CHD4"     "SUZ12"    "CDC6"     "CHD3"     "ORC1"
#>  [7] "FUS"      "MCM4"     "AURKB"    "MRPS14"   "RPL26L1"  "RPS13"
#> [13] "RPS23"    "BRD2"     "CHD5"     "KDM2A"    "SUV420H1" "SUV420H2"
#> [19] "KAT7"     "SETD8"    "MSH3"     "ERCC3"    "MARCKS"   "CDCA2"
#> [25] "CDCA5"    "CDCA7L"   "ATAD2B"   "NUDT21"   "KIF22"    "RPS25"
```

```
#> [31] "VDAC3"     "CPSF7"     "DIEXF"     "POLR1E"    "XIRP2"     "ZFP91"
#> [37] "ARHGAP11A" "CCDC137"   "KLHL15"    "OR10G3"
```

```
terms
#> [1] "Acidic Patch" "Chromatin"    "Nucleosome"
```

We can now run DeSciDe on this list in the most simple form. This will produce our figures and our data table of results.

```
results <- descide(genes_list = genes, terms_list = terms)
```

##### Results Expected from DeSciDe

```
#> # A tibble: 6 x 14
#>   Gene     `Acidic Patch` Chromatin Nucleosome Total PubMed_Rank Degree
#>   <chr>             <int>     <int>      <int> <dbl>       <int>  <dbl>
#> 1 BRD4                  2       718         29   749           1     14
#> 2 SUV420H1              2        39          8    49           2      4
#> 3 CHD4                  1       202        131   334           3      8
#> 4 KDM2A                 1        49          5    55           4      6
#> 5 SUZ12                 0       237         13   250           5      5
#> 6 CDC6                  0       191          6   197           6     10
#> # i 7 more variables: Clustering_Coefficient_Percent <dbl>,
#> #   Clustering_Coefficient_Fraction <chr>, Connected_Component_id <dbl>,
#> #   Nodes_in_Connected_Component <dbl>,
#> #   total_number_of_connected_components <dbl>, Connectivity_Rank <int>,
#> #   Category <chr>
```

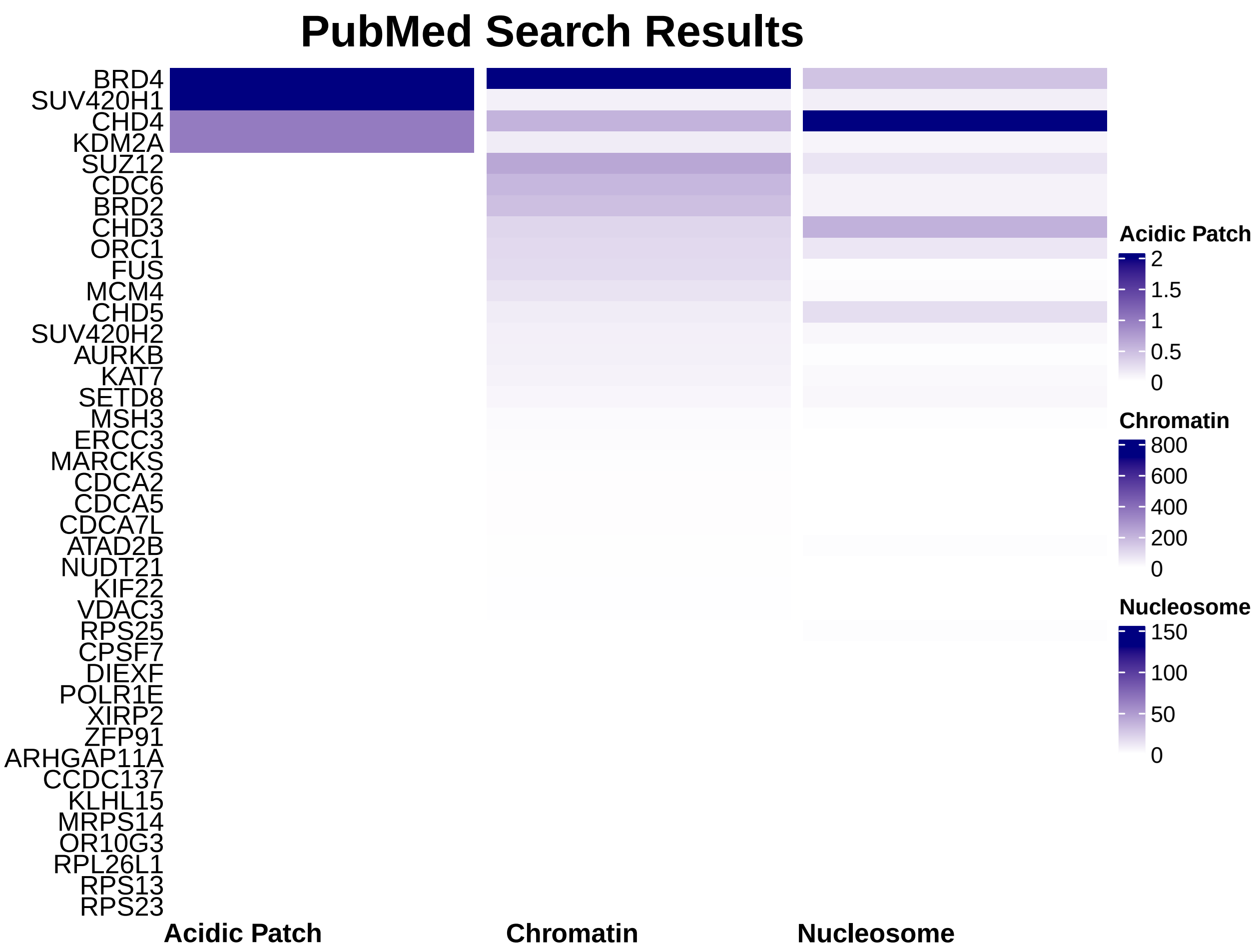
PubMed Search Results
BRD4
SUV420H1
CHD4
KDM2A
SUZ12
CDC6
BRD2
CHD3
ORC1
FUS
MCM4
CHD5
SUV420H2
AURKB
KAT7
SETD8
MSH3
ERCC3
MARCKS
CDCA2
CDCA5
CDCA7L
ATAD2B
NUDT21
KIF22
VDAC3
RPS25
CPSF7
DIEXF
POLR1E
XIRP2
ZFP91
ARHGAP11A
CCDC137
KLHL15
MRPS14
OR10G3
RPL26L1
RPS13
RPS23
Acidic Patch
Chromatin
Nucleosome
Acidic Patch
2
1.5
1
0.5
0
Chromatin
800
600
400
200
0
Nucleosome
150
100
50
0

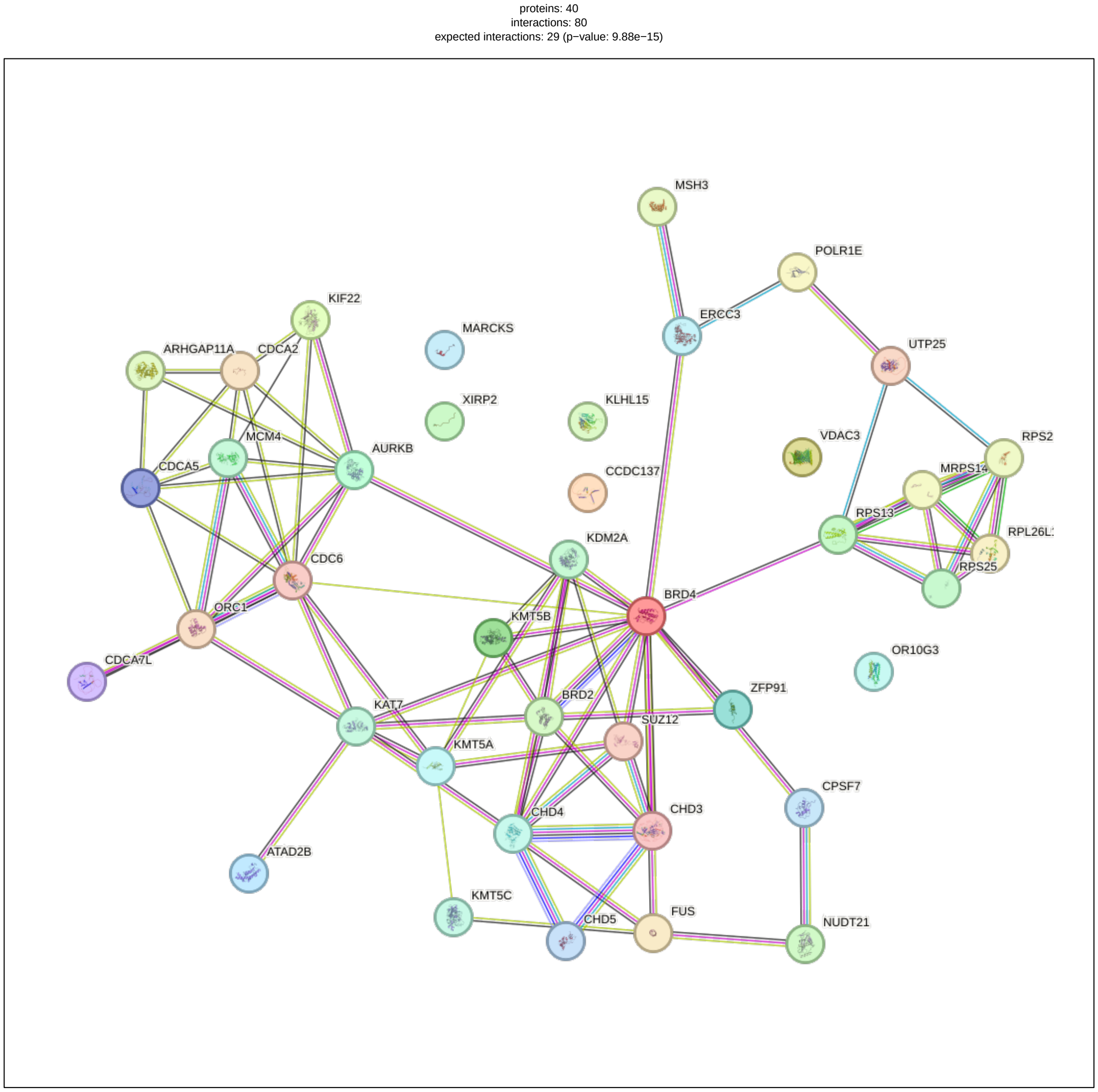

```
#> 2025-04-30 10:47:01.740171: Inside plot_clustering function
#> 2025-04-30 10:47:01.740365: After conversion - Data types of columns:
#> 'data.frame':    40 obs. of  8 variables:
#>  $ Gene_Symbol                       : chr  "BRD4" "CDC6" "AURKB" "CHD4" ...
#>  $ Degree                            : num  14 10 8 8 7 7 6 6 6 6 ...
#>  $ Clustering_Coefficient_Percent    : num  17.6 37.8 53.6 42.9 47.6 ...
#>  $ Clustering_Coefficient_Fraction   : chr  "16/91" "17/45" "15/28" "12/28" ...
#>  $ Connected_Component_id            : num  1 1 1 1 1 1 1 1 1 1 ...
#>  $ Nodes_in_Connected_Component      : num  34 34 34 34 34 34 34 34 34 34 ...
#>  $ total_number_of_connected_components: num  7 7 7 7 7 7 7 7 7 7 ...
#>  $ Connectivity_Rank                 : int  1 2 3 4 5 6 7 8 9 10 ...
#> NULL
```

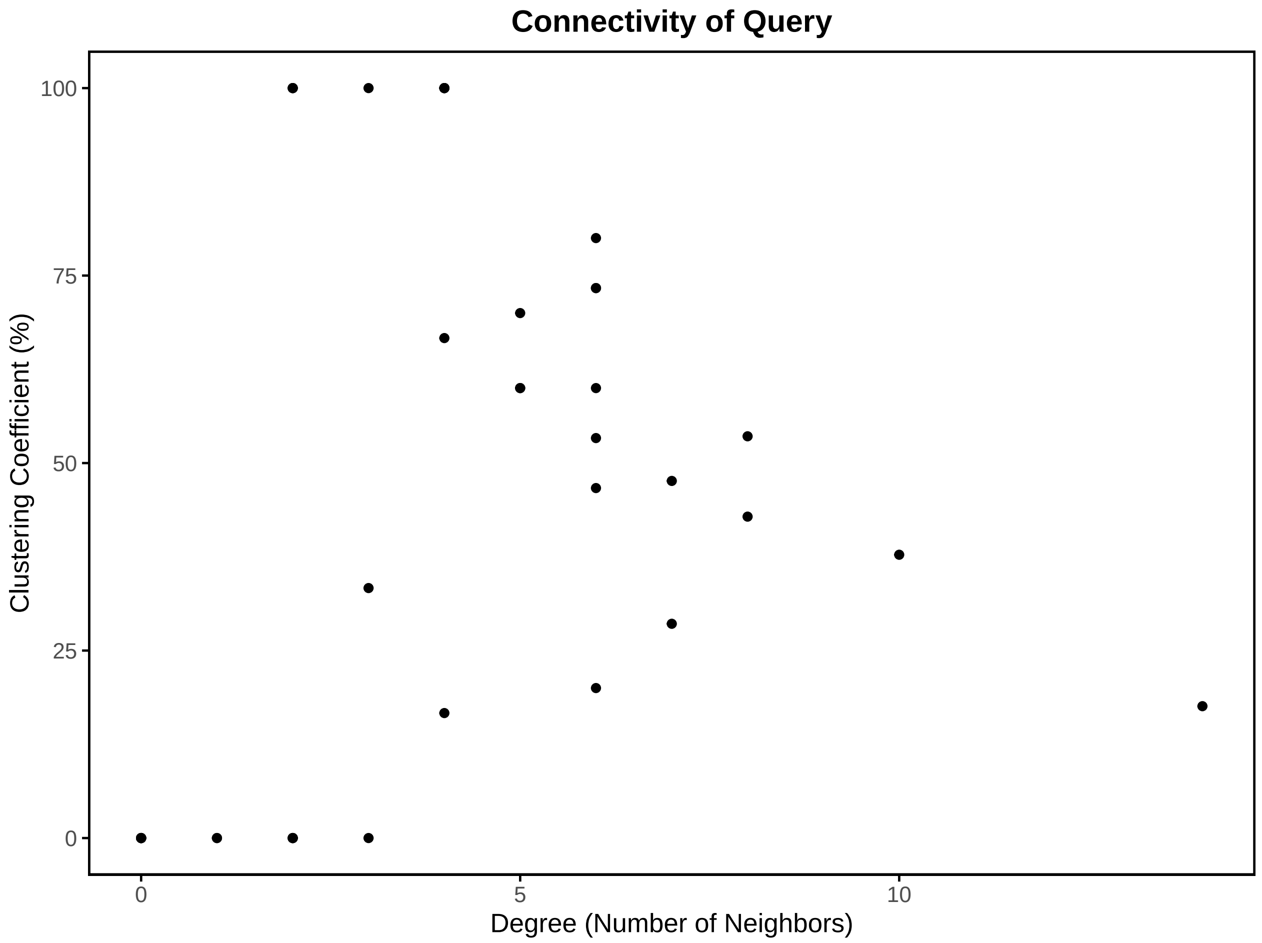
Connectivity of Query
Clustering Coefficient (%)
100
75
50
25
0
0
5
10
Degree (Number of Neighbors)

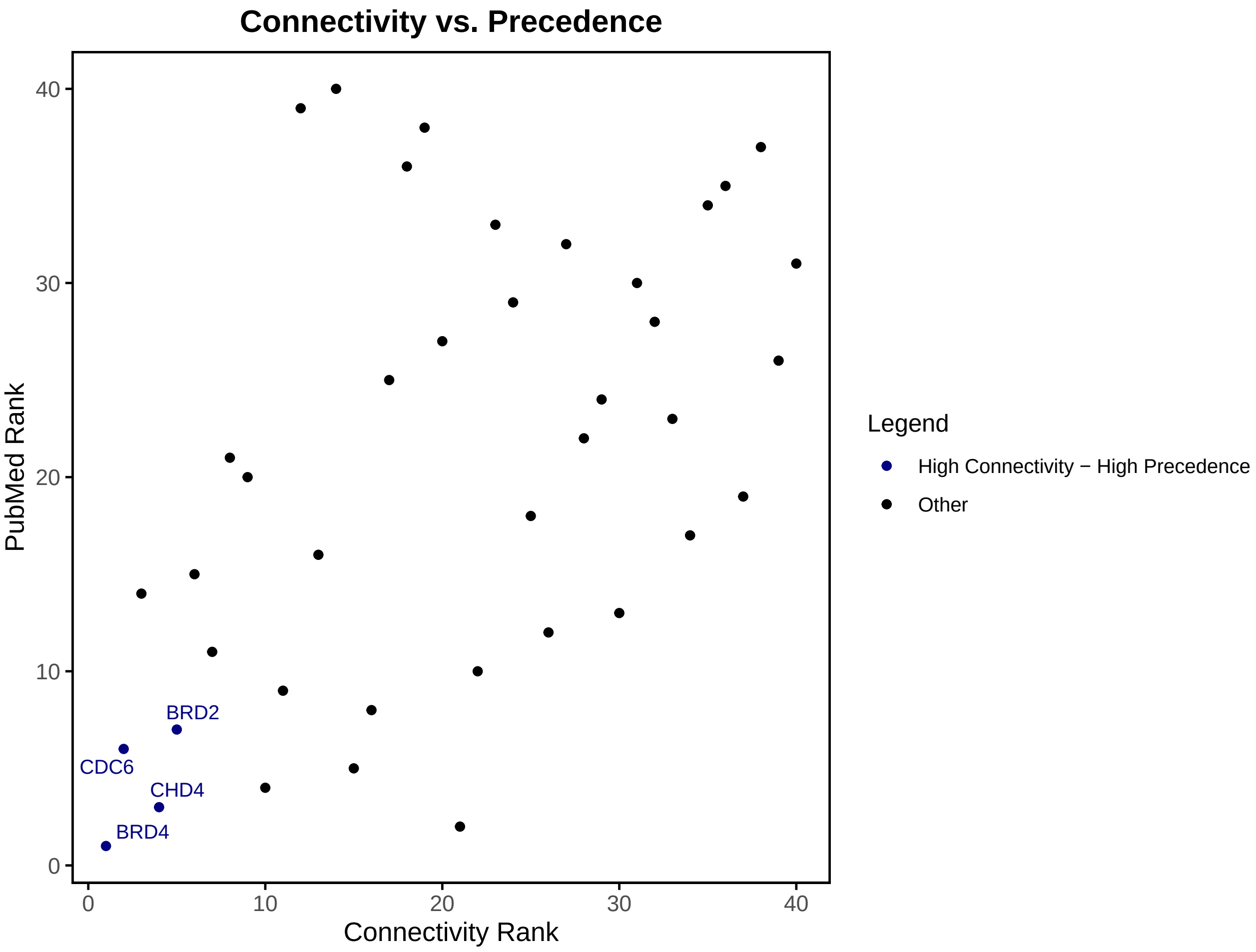

#### Modifications to DeSciDe Search

##### PubMed Ranking Method: Important

By default, the PubMed search results are sorted for ranking in order of the terms provided. Here we gave "Acidic Patch" first, so the table is sorted with preference for "Acidic Patch" and then sorted by "Chromatin" and finally by "Nucleosome". You can see that this results in CHD4 being ranked as lower precedence that SUV420H1 even though it has more publications in both the chromatin and nucleosome searches. You must be aware of this when creating your search term list. This weighting has been incorporated to help users emphasize terms highly specific to their research that may have significantly lower number of results than a broader term incorporated into the search (i.e. Chromatin in our search).

If a user does not want to weight their search to a specific term, they can use the argument `rank_method = "total"` to rank the genes by the sum of publication numbers across each search term.

```
results_total <- descide(genes_list = genes, terms_list = terms, rank_method = "total")
```

Observe the changes in the PubMed rankings and the Heatmap as a result of weighting by "total". SUV420H1 is dropped from rank 2 down to 13 as a result of the change in weighting. It is important for users to be aware of the differences in these two methods. Both methods, weighted or total, can provide valuable insight into a data set, but the user needs to be conscientious of which method is going to best serve their purpose.

```
head(results$summary_results)
#> # A tibble: 6 x 14
#>   Gene     `Acidic Patch` Chromatin Nucleosome Total PubMed_Rank Degree
#>   <chr>             <int>     <int>      <int> <dbl>       <int>  <dbl>
```

```
#> 1 BRD4                  2       718          29   749           1      14
#> 2 SUV420H1              2        39           8    49           2       4
#> 3 CHD4                  1       202         131   334           3       8
#> 4 KDM2A                 1        49           5    55           4       6
#> 5 SUZ12                 0       237          13   250           5       5
#> 6 CDC6                  0       191           6   197           6      10
#> # i 7 more variables: Clustering_Coefficient_Percent <dbl>,
#> #   Clustering_Coefficient_Fraction <chr>, Connected_Component_id <dbl>,
#> #   Nodes_in_Connected_Component <dbl>,
#> #   total_number_of_connected_components <dbl>, Connectivity_Rank <int>,
#> #   Category <chr>
```

```
head(results_total$summary_results)
#> # A tibble: 6 x 14
#>   Gene   `Acidic Patch` Chromatin Nucleosome Total PubMed_Rank Degree
#>   <chr>           <int>     <int>      <int> <dbl>       <int>  <dbl>
#> 1 BRD4                2       718         29   749           1     14
#> 2 CHD4                1       202        131   334           2      8
#> 3 SUZ12               0       237         13   250           3      5
#> 4 CDC6                0       191          6   197           4     10
#> 5 BRD2                0       169          6   175           5      7
#> 6 CHD3                0       109         38   147           6      5
#> # i 7 more variables: Clustering_Coefficient_Percent <dbl>,
#> #   Clustering_Coefficient_Fraction <chr>, Connected_Component_id <dbl>,
#> #   Nodes_in_Connected_Component <dbl>,
#> #   total_number_of_connected_components <dbl>, Connectivity_Rank <int>,
#> #   Category <chr>
```

The more specific and niche the search terms are, the more likely you will want to use weighted. In the example here, "Acidic Patch" is a specific term that does not appear in many publications in combination with the provided gene list. The experiment associated with this dataset was studying mutations to the acidic patch of histone H2A, so as a primary variable in the experiment it is valuable to weight the results to this term, to ensure that the variables of the system (WT or mutants of histone H2A acidic patch) would be represented in DeSciDe's results. Searching by total, however, can provide a good starting place for analysis when a user is not sure what terms to prioritize or does not yet know what precedence there is among their gene list for the terms they include.

##### Modifications to STRING Search

When searching the STRING database for gene interactions, there are a variety of variables that can be adjusted. These variable modifications have been included in DeSciDe's usage. First is the specification of the species of interest. The package defaults to search for human genes, but this can be changed by the user with argument `species =` as shown below:

```
# Change species to mus musculus for STRING search.
descide(genes_list = genes, terms_list = terms, species = 10090)
```

Additionally, STRING creates scores for each interaction that occurs within a network. This score can be adjusted to increase or decrease the confidence in interactions. High confidence interactions have a score of 1000 and low confidence interactions have a score of 0. The default score for STRING is 400, which is the default used within DeSciDe. To change the STRING score minimum value, the user can use the argument `score_threshold =`:

```r
# Change STRING score threshold to 600.
descide(genes_list = genes, terms_list = terms, score_threshold = 600)
```

The last variable that can be changed within DeSciDe to modify the STRING search is modifying the network type. By default DeSciDe uses a full network search. The full network includes all functional relationships between genes/proteins whether they directly interact or are related to each other for other reasons such as homology, gene ontology, etc. The other option is to limit the network to explicitly physical interactions. This parameter can be changed within DeSciDe by using the argument `network_type =`:

```r
# Change STRING network type to only include physical interactions.
descide(genes_list = genes, terms_list = terms, network_type = "physical")
```

We can run just the STRING search to see the differences in the network produced by using full or physical network. With the full network, we see many more interactions between genes:

```r
# Run STRING search and display network with full network.
full_string <- search_string_db(genes_list = genes, network_type = "full")
plot_string_network(full_string$string_db, full_string$string_ids)
```

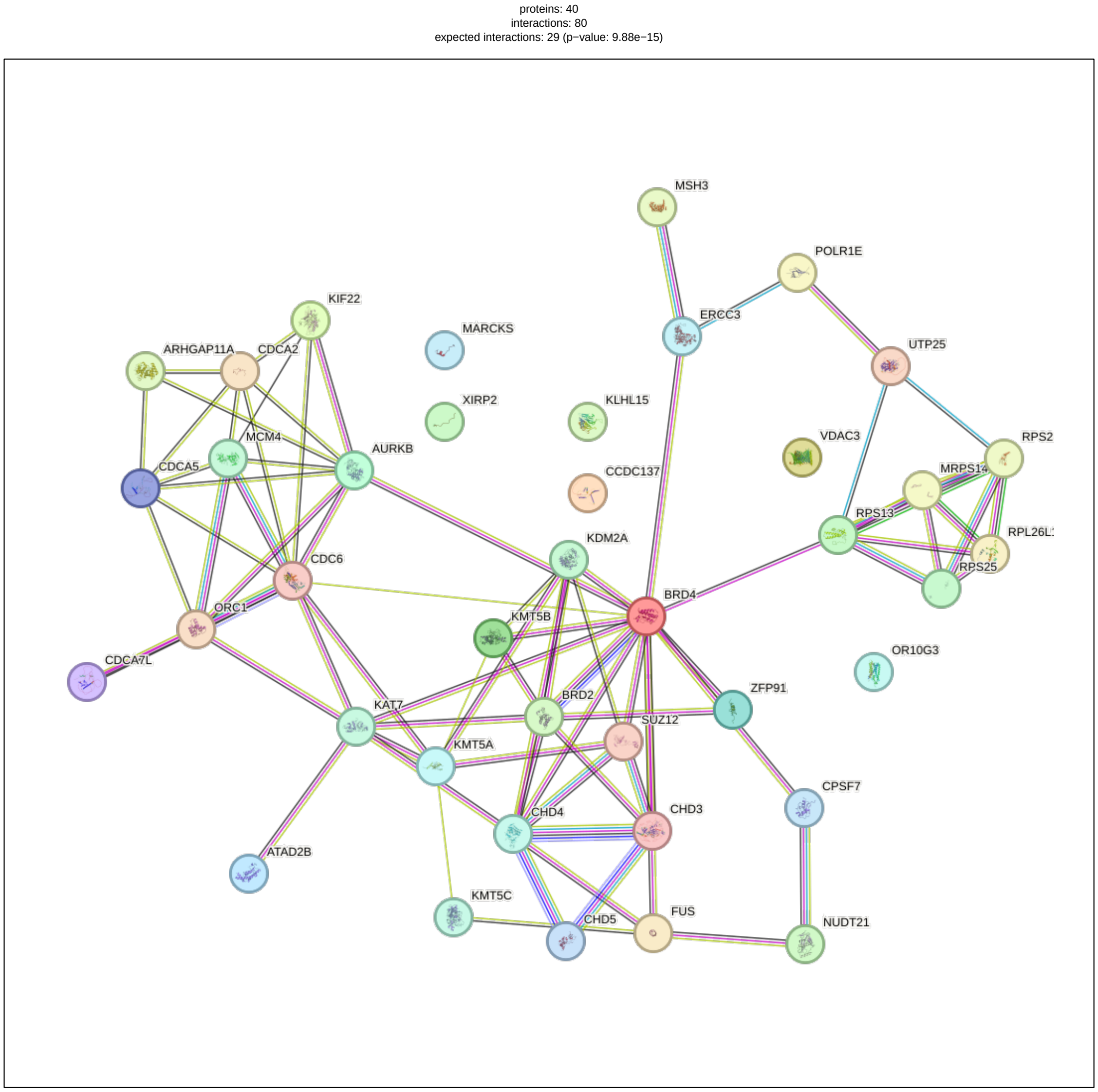

With the more stringent physical network, we see fewer interactions.

```r
# Run STRING search and display network with physical network.
physical_string <- search_string_db(genes_list = genes, network_type = "physical")
plot_string_network(physical_string$string_db, physical_string$string_ids)
```

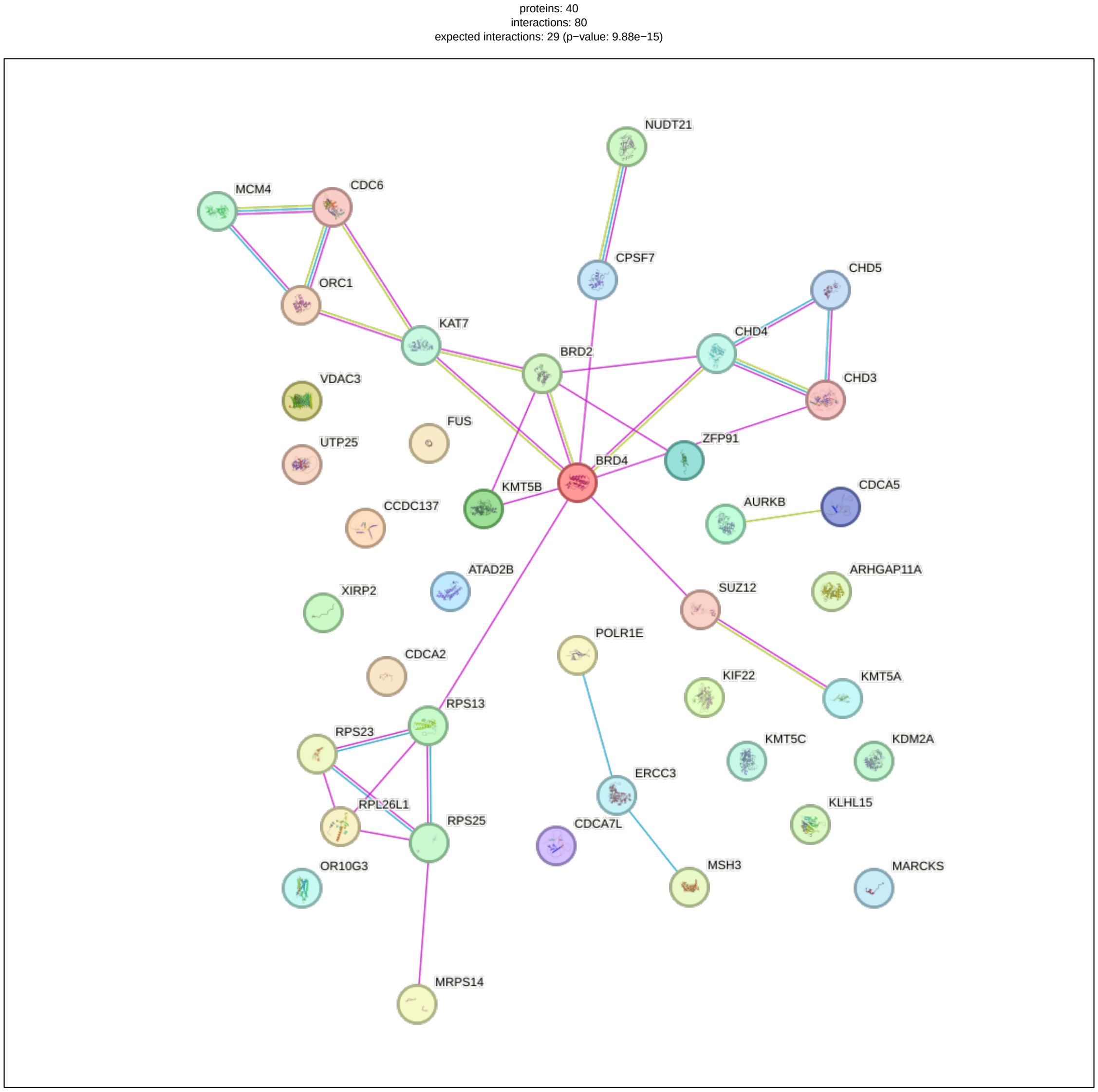

#### Modifications to DeSciDe Classifications

The rank list of PubMed and STRING results are combined into a summary file. These ranks are then used to classify the genes as either high connectivity - high precedence (high confidence genes) or high connectivity - low precedence (novel genes). The classification is conducted by setting thresholds based on the length of the gene list. The default threshold is 20%. High connectivity - high precedence genes are those that fall in the 20th percentile of both pubmed and string ranks (i.e. for a list of 100 genes, these are genes ranked 1-20). High connectivity - low precedence genes are those that rank in the 20th percentile for STRING rank and the 80-100th percentile for PubMed results (i.e. ranks 81-100 in a list of 100 genes). This value can be adjusted by the user to make the classification more or less stringent by using the argument `threshold_percentage =`.

To change the threshold percentage in the complete DeSciDe pipeline:

```
# Command to adjust threshold_percentage for full descide pipeline.
results <- descide(genes_list = genes, terms_list = terms, threshold_percentage = 50)
```

We can compare the 20% and 50% threshold results by running just the `combine_summary()` and `plot_connectivity_precedence()` functions:

```
# Calculate and plot threshold of 20%.
threshold_20 <- combine_summary(pubmed_search_results = results$pubmed_results,
                  string_results = results$string_results, threshold_percentage = 20)
plot_connectivity_precedence(combined_summary = threshold_20)
```

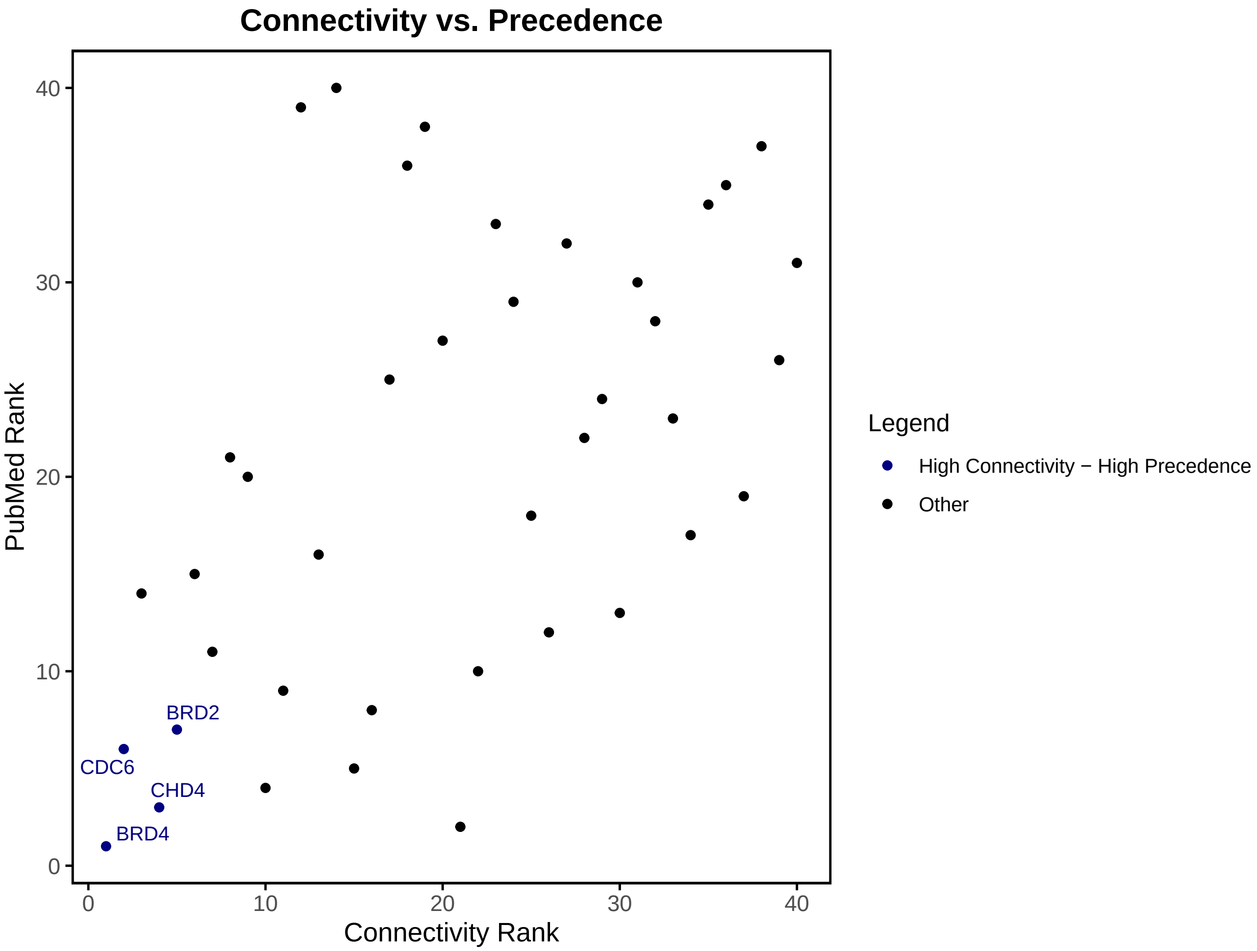

```
# Calculate and plot threshold of 50%.
threshold_50 <- combine_summary(pubmed_search_results = results$pubmed_results,
                  string_results = results$string_results, threshold_percentage = 50)
plot_connectivity_precedence(combined_summary = threshold_50)
```

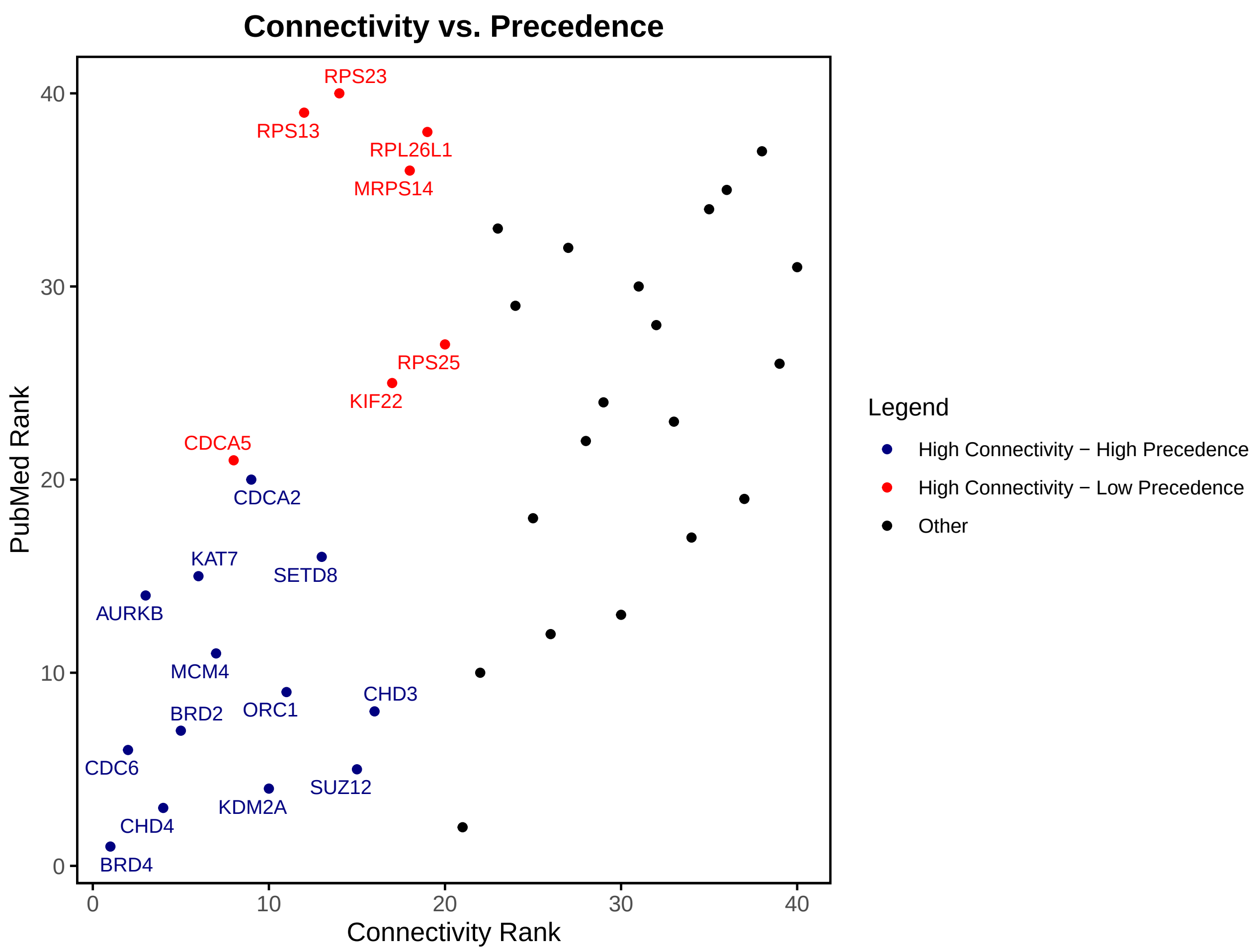

```
head(threshold_20)
#> # A tibble: 6 x 14
#>   Gene     `Acidic Patch` Chromatin Nucleosome Total PubMed_Rank Degree
#>   <chr>             <int>     <int>      <int> <dbl>       <int>  <dbl>
#> 1 BRD4                  2       718         29   749           1     14
#> 2 SUV420H1              2        39          8    49           2      4
#> 3 CHD4                  1       202        131   334           3      8
#> 4 KDM2A                 1        49          5    55           4      6
#> 5 SUZ12                 0       237         13   250           5      5
#> 6 CDC6                  0       191          6   197           6     10
#> # i 7 more variables: Clustering_Coefficient_Percent <dbl>,
#> #   Clustering_Coefficient_Fraction <chr>, Connected_Component_id <dbl>,
#> #   Nodes_in_Connected_Component <dbl>,
#> #   total_number_of_connected_components <dbl>, Connectivity_Rank <int>,
#> #   Category <chr>
head(threshold_50)
#> # A tibble: 6 x 14
#>   Gene     `Acidic Patch` Chromatin Nucleosome Total PubMed_Rank Degree
#>   <chr>             <int>     <int>      <int> <dbl>       <int>  <dbl>
#> 1 BRD4                  2       718         29   749           1     14
#> 2 SUV420H1              2        39          8    49           2      4
#> 3 CHD4                  1       202        131   334           3      8
#> 4 KDM2A                 1        49          5    55           4      6
#> 5 SUZ12                 0       237         13   250           5      5
```

```
#> 6 CDC6                    0       191          6   197            6      10
#> # i 7 more variables: Clustering_Coefficient_Percent <dbl>,
#> #   Clustering_Coefficient_Fraction <chr>, Connected_Component_id <dbl>,
#> #   Nodes_in_Connected_Component <dbl>,
#> #   total_number_of_connected_components <dbl>, Connectivity_Rank <int>,
#> #   Category <chr>
```

This change to threshold percentage just modifies the annotation of genes in the category column of the summary results and the annotations of the genes on the scatter plot of connectivity vs precedence. Using higher threshold can help get a better visual idea of where more genes fall in the analysis, but lowering the threshold can help narrow in on select genes to conduct follow up experiments on.

###### Exporting Tables and Graphs

For ease of use, we have incorporated a feature to easily export all of the tables and graphs to the users desired destination. To do this, you can use the arguments `export = TRUE` and `file_directory = "your/desired/directory"` to export to your desired directory. Furthermore, you can specify the format of the tables using `export_format =`, which you can specify as "csv", "tsv", or "excel". Here is an example of running the full DeSciDe pipeline to be exported:

```
# Code to run DeSciDe and export all plots and tables to desired directory.
descide(genes_list = genes, terms_list = terms, export = TRUE,
        file_directory = "your/desired/directory", export_format = "excel")
```

#### All Functions Available for DeSciDe

Each step of DeSciDe can be run individually if you wish to do so. Below we briefly list each function and all of their arguments. To see more information, you can use the help function in R studio to see the R documentation for each function (i.e. `?descide` or `?plot_connectivity_precedence`)

##### Function to run entire pipeline

```
descide(
  genes_list,
  terms_list,
  rank_method = "weighted",
  species = 9606,
  network_type = "full",
  score_threshold = 400,
  threshold_percentage = 20,
  export = FALSE,
  file_directory = NULL,
  export_format = "csv"
)
```

##### Function to run PubMed search

```
search_pubmed(genes_list, terms_list, rank_method = "weighted")
```

**Function to plot heatmap**

```
plot_heatmap(pubmed_search_results, file_directory = NULL, export = FALSE)
```

**Function to search STRING**

```
search_string_db(
  genes_list,
  species = 9606,
  network_type = "full",
  score_threshold = 400
)
```

**Function to plot STRING network**

```
plot_string_network(
  string_db,
  string_ids,
  file_directory = NULL,
  export = FALSE
)
```

**Function to plot STRING clustering metrics**

```
plot_clustering(string_results, file_directory = NULL, export = FALSE)
```

**Function to create summary file and classify genes based on ranks.**

```
combine_summary(
  pubmed_search_results,
  string_results,
  file_directory = NULL,
  export_format = "csv",
  export = FALSE,
  threshold_percentage = 20
)
```

**Function to plot connectivity vs. precedence.**

```
plot_connectivity_precedence(
  combined_summary,
  file_directory = NULL,
  export = FALSE
)
```
